# Multi-Locus Molecular Phylogeny of *Solenopsis* (Hymenoptera: Formicidae)

**DOI:** 10.1101/2020.06.05.136945

**Authors:** Scott Shreve, Rafael Achury, Kevin Johnson, Andrew Suarez, Dietrich Gotzek

## Abstract

The myrmicine ant genus *Solenopsis* is species-rich, globally distributed, and is often a common and ecologically important faunal element of the leaf litter. The genus is also well-known for containing several widely distributed tramp species and some of the worst invasive species in the World (the Red Imported Fire Ant, *S. invicta*, and the Tropical Fire Ant, *S. geminata*). Although not hyper-diverse and despite its ecological and economic importance, *Solenopsis* has long frustrated systematists due its lack of reliable diagnostic characters and no phylogenetic hypothesis exists to date. We present a preliminary multi-locus molecular phylogenetic analysis of *Solenopsis* to address this knowledge gap. Our analyses recover mostly well-supported phylogenetic hypothesis, which suggests *Solenopsis* arose in the Neotropics and spread to all other continents (except Antarctica). Importantly, it demonstrates problems with current systematic understanding of the genus, but provides an evolutionary framework upon which to build future research.

## Introduction

Ants are important elements in most terrestrial ecosystems due to both their significant contribution to the animal biomass in an ecosystem and their ability to alter ecosystems (Folgarait 1998). Many ant species have spread far beyond to their native ranges, in some cases achieving worldwide distributions, as a result of their associations with humans. It has been estimated that approximately 150 species (McGlynn 1999) fall into this group, but the number of non-native species has undoubtedly grown since then, both as a result of new knowledge of ant distributions coming to light and of new movements of previously localized species by increased human activity. These widespread ants with many non-native populations can be treated as two distinct categories (Holway et al. 2002). The larger category, often termed “tramp ants” rely primarily on human-mediated dispersal and tend to maintain a close association with humans (Hölldobler & Wilson 1990; Passera 1994). “Invasive ants,” on the other hand, once they have reached new areas via human activity, are able to break the association with humans and penetrate natural ecosystems (Holway et al. 2002). The effect of tramp ants is largely urban or agricultural in nature, whereas invasive ants can also reduce native ant diversity and adversely affect other organisms dependent on the native ant community (Holway et al. 2002). Invasive ants may also aid in the dispersal of other invasive arthropods; 29 species of potentially phoretic arthropods were collected off of alates from nests at a single locality in the southeastern United States (Moser & Blomquist 2011).

Two of the six most widespread, invasive ant species are in the genus *Solenopsis*: *S. invicta* and *S. geminata* (Holway et al. 2002). These two species are unusual among invasive ants, however, in that dispersal is often accomplished by winged flight of the female reproductive forms (DeHeer et al. 1998; Holway et al. 2002). If winged dispersal is phylogenetically conserved, other species of *Solenopsis* ants may be pre-adapted to become invasive species, or at the very least have facilitated the spread of members of the genus throughout the world.

The genus *Solenopsis* has historically been classified into three primary groups on the basis of natural history: fire ants, thief ants, and social inquiline parasites (Ettershank 1966). However, only the fire ants form a well-defined group, and the inquiline social parasites have likely arisen multiple times within the genus (Trager 1991; Pitts et al. 2005). The 23 species of fire ants are currently classified into four species groups (Table 1; Trager 1991; Pitts et al. 2005, 2018). The final category of *Solenopsis* ants are the thief ants, so called because they tend to nest near to or inside other ant colonies and steal their brood (Pacheco & Mackay 2013). Globally, there are currently 171 described species of thief ants (Bolton 2016) organized into eleven species complexes (Table 1; Pacheco & Mackay 2013; Galkowski et al. 2010). However, there has been no taxonomic or phylogenetic treatment of the entire genus, and the phylogenetic status and relationships within and among the species groups are unknown. In addition to the 196 recognized *Solenopsis* species (Bolton 2016), there are likely a large number of undescribed species, especially in the taxonomically neglected thief ants. Recent molecular species delimitation studies suggest that cryptic species are common even among the relatively well-studied fire ants (Ross & Shoemaker 2005; Ross et al. 2007, 2010; Chialvo et al. 2017) and can thus be expected in the genus as a whole.

**Table 1.** Recognized species complexes of the genus *Solenopsis*. Fire ant classification follows Pitts et al. (2005, 2018) and thief ant classification follows Pacheco & Mackay (2013) for New World (NW) species and Galkowski et al. (2010) for Old World (OW) species. Greater equal signs indicate the presence of undescribed species or geographically restricted tallying.

| Species Complex | Number of Species | Distribution |
| --- | --- | --- |
| “Fire Ants” |  |  |
| *geminata* | 4 | NW (OW) |
| *saevissima* | ≥16 | NW (OW) |
| *tridens* | 2 | NW |
| *virulens* | 1 | NW |
| “Thief Ants” |  |  |
| *brevicornis* | 3 | NW |
| *debilior* | ≥1 | OW |
| *fugax* | ≥14 | NW+OW |
| *globularia* | >4 | NW |
| *lusitanica* | ≥4 | OW |
| *molesta* | >34 | NW |
| *nigella* | >10 | NW |
| *orbula* | ≥25 | OW |
| *pygmaea* | 12 | NW |
| *stricta* | >2 | NW |
| *wasmannii* | 4 | NW |

In the present study, we use molecular sequence data of five genes to generate a phylogenetic hypothesis for the relationships among species in the genus *Solenopsis*. We use this phylogeny to determine if robust species groups can be identified and if these species groups fall out into meaningful biogeographic clusters. We also examine where within the genus fire ants originate. Finally, we use our diverse sampling to examine the distribution of introduced species across the phylogeny and use these results to correct taxonomic uncertainty for widespread tramp species.

## Methods

### Sample Collection

Species identifications and collection localities for specimens are given in Table 2. Samples were obtained from various natural history museums and private collections. When possible, specimens were identified to species or species complex (Pacheco & Mackay 2013; Galkowski et al. 2010). Many specimens in our dataset are either undescribed species or could not unequivocally be identified to species due to lack of workable keys, especially in the Old World. In these cases, specimens were assigned to species complexes or were assigned country codes (e.g., CAM-1 to CAM-3). We included nuclear gene data of all Solenopsidini from Ward et al. (2015) to complement our own outgroup sampling.

**Table 2.**
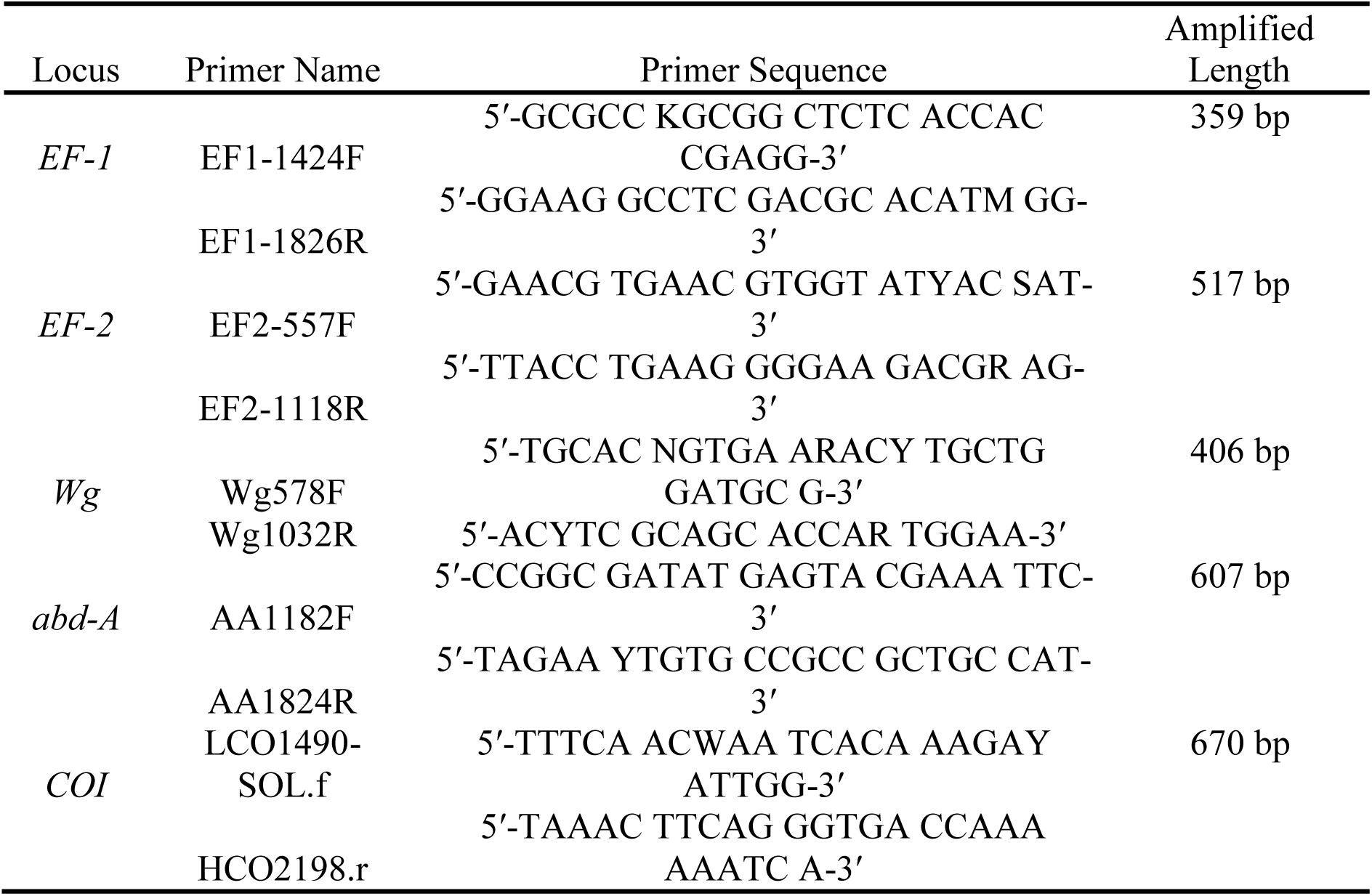
Primer sequences used in PCR amplification, amplified fragment length, and nucleotide substitution models used in ML analysis. Lengths of each gene are aligned lengths after primer sequences and un-alignable regions were removed.

### DNA Amplification and Sequencing

The phylogenetic relationships among the *Solenopsis* genus were inferred using four nuclear genes and one mitochondrial gene, totaling 2,567 aligned bp. Primer sequences and sequence lengths for the five genes used in this study are given in Table 2. All PCR reactions were performed in 12.5 µL using Taq-Pro (Denville Scientific Inc.). The final reagent concentrations for the *EF-1a* amplifications were 1.0 µL of extracted DNA in 1.1X Taq-Pro 2.0 mM Mg^2+^ buffer, 0.4 µM of each primer, and 0.08 mg/mL bovine serum albumin (BSA). We used a touchdown PCR procedure, with an initial denaturing step of 95°C for 2 min; followed by 95°C for 30 s, 62°C for 30 s, and 72°C for 30 s, with the annealing temperature decreasing by 0.4°C/cycle for 12 cycles; 95°C for 30 s, 57°C for 30 s, and 72°C for 30 s for an additional 30 cycles, and a final elongation step of 72°C for 5 min. The final reagent concentrations for the *EF-2* were 1.0 µL of extracted DNA in 1.1X Taq-Pro 1.5 mM Mg^2+^ buffer, 0.4 µM of each primer, and 0.06 mg/mL BSA. We used a similar touchdown PCR procedure, differing from that employed in *EF-1* changing the starting annealing temperature at 60°C, then decreasing by 0.4°C/cycle for 10 cycles, and finally settling at 56°C for 30 cycles.

PCR amplifications of *wingless* (*Wg*) used final reagent concentrations of 1.0 µL of extracted DNA, 1.1X Taq-Pro 2.0 mM Mg^2+^ buffer, 0.35 µM of each primer, and 0.08 mg/mL BSA. The touchdown procedure began with an annealing temperature of 62°C, decreasing by 0.4°C/cycle for 10, and ending at 58°C for 30 cycles. *Abdominal-A* (*Abd-A*) amplifications used 1.25 µL extracted DNA in 1.0X Taq-Pro 2.0 mM Mg^2+^ buffer, 0.4 µM of each primer, and 0.08 mg/mL BSA. The reaction conditions were identical to those used with EF-1. PCR amplifications of COI used final reagent concentrations of 1.0 µL of extracted DNA, 1.2X Taq-Pro 1.5 mM Mg^2+^ buffer, 0.4 µM of each primer, and 0.08 mg/mL BSA. Reaction conditions were 95°C for 2 min; 95°C for 30 s, 45°C for 30 s, and 72°C for 40 s for 10 cycles; 95°C for 30 s, 51°C for 30 s, and 72°C for 40 s; and 72°C for 5 min.

All sequences were deposited in GenBank under accession numbers MT550038 – MT550618.

### Phylogenetic Analyses

Alignment of sequences for each locus was conducted using *MAFFT 6* (Katoh et al. 2002; Katoh & Toh 2008) using the E-INS-I algorithm, and alignments were adjusted by eye. The dataset was partitioned into 16 regions corresponding to the three codon positions of each of the five genes sequenced and the intron region of *COI*. Optimal partitioning and nucleotide substitution models for each partition were chosen using *PartitionFinder 1.1.1* (Lanfear et al. 2012) implementing linked branch lengths, BIC for model selection, and greedy search scheme of all models. Models selected for each partition are given in Table 3.

**Table 3.** PartitionFinder results for 16 partitions using BIC model selection of 24 nucleotide substition models with greedy search option and linked branch lengths.

| Partition | Best Model | Subset Partitions |
| --- | --- | --- |
| p1 | K80+I+G | EF1 c1, EF2 c2 |
| p2 | GTR+G | EF1 c2, Wg c3 |
| p3 | GTR+I+G | abdA c1, COI intron, EF1 c3, EF2 c1, Wg c2 |
| p4 | SYM+I+G | EF2 c3 |
| p5 | JC | Wg c1 |
| p6 | F81+I+G | abdA c2 |
| p7 | GTR+G | abdA c3 |
| p8 | GTR+I+G | COI c3 |
| p9 | SYM+G | COI c1 |
| p10 | GTR+I+G | COI c2 |

Maximum likelihood analysis (ML) using partitions by genes was conducted with twenty independent runs in *GARLI 2.1* (Zwickl 2006). Branch support was determined with a bootstrap analysis consisting of 1,000 replicates, also implemented in *GARLI*, and a Bayesian-like transformation of the approximate likelihood ration test (aBayes; Anisimova et al. 2011) implemented in IQTREE (-m MFP+MERGE -alrt 1000 -abayes). A Bayesian phylogenetic analysis (BI) was carried out using *MrBayes v3.2.5* (Huelsenbeck & Ronquist 2001; Ronquist & Huelsenbeck & Ronquist 2003). Three independent analyses were run for 5×10^7^ generations, sampling every 5,000 generations. Each run consisted of one cold and three heated chains. The temperature parameter in *MrBayes* was decreased to 0.008 to ensure sufficient swapping among the chains. An identical analysis was conducted without data to assess potential undue influence of the priors. Convergence among chains was assessed using typical convergence diagnostics and metrics implemented in *MrBayes, Tracer 1.6, RWTY 1.0.1* (Warren et al. 2017) (an *R 3.4.1* [R Core Team 2017] implementation and extension of *AWTY* [Nylander et al. 2008a]), and *coda 0.18-1* (Plummer et al. 2006).

We also visualized tree space using multi-dimensional scaling (MDS; Hillis et al. 2002) implemented in *RWTY* and the *R* package *Treespace 1.0.0* (Jombart et al. 2017). We compared Robinson – Foulds symmetric distances (Robinson & Foulds 1981), which are appropriate measures for comparing tree topologies (Kuhner & Yamato 2014).

### Biogeography of Solenopsis

Sampled *Solenopsis* were assigned to one of ten biogeographic realms (*sensu* Holt et al. 2013) based on collection locality: (A) Afrotropical, (B) Australian, (C) Madagascan, (D) Nearctic, (E) Neotropical, (F) Oceanian, (G) Oriental, (H) Palearctic, (I) Panamanian, and (J) Saharo-Arabian. All outgroup taxa are Neotropical in distribution, except for *Austromorium* and *Monomorium antarcticum*, which occur in the Australian realm (Ward et al. 2015). The ancestral biogeographic distributions of *Solenopsis* were inferred by statistical dispersal-vicariance analysis (S-DIVA; Ronquist 1997; Nylander et al. 2008b) and Bayesian binary MCMC (BBM;) implemented in *RASP 3.2* (Yu et al. 2015; Ali et al. 2012). For both analyses, we limited the maximum number of regions of reconstructed ranges to four. The BBM analysis consisted of 50 independent MCMC chains, each with 5,000,000 generations sampled every 1,000 generations with a burnin of 2,000 generations. We used the F81+G (unequal rates with rate heterogeneity) with null root distribution was used as the underlying model and all calculations were based on the fully bifurcating ML topology. For the S-DIVA analysis, we randomly sampled 1,000 trees from the 15,000 post-burnin trees from the BI analysis to account for phylogenetic uncertainty. Since our outgroups are not representative of the diversity of biogeographic distributions of their higher-level taxa, we also replicated the analyses with the outgroups excluded to determine the extent to which the inferences depended on our outgroup choice.

## Results

### DNA Sequencing

Of the 216 *Solenopsis* and outgroup samples attempted, 124 samples had sequences for at least three of the five genes, with 69 of the samples having all five genes. To this we added four samples of particular taxonomic or biogeographic interest (the samples from Azerbaijan and Borneo, *S. metanotali*, and *S. wasmanni*) that only had sequences for two loci. This resulted in a final dataset of 128 operational taxonomic units (OTUs). All but two of the samples in the final dataset possessed *Wg* sequences. All four added samples with two genes had *Wg* as one of the two sequenced loci in order to facilitate the accurate placement of these samples in the phylogeny.

### Phylogenetic Analysis

The phylogenetic analyses recovered congruent and well resolved topologies (Figure 1). However, our dataset is not optimally informative at the deeper levels of the phylogeny, leading to poorly supported deeper nodes (Table 4) and difficulties to infer the evolutionary history of the outgroups (not shown). For some of the weaker supported nodes, the level of branch support also differed depending on measure of branch support (Table 4). However, both the ML and BI analyses most strongly support a monophyletic genus and the topology within *Solenopsis* is completely concordant with that of Ward et al. (2015). We thus recognized and discuss five well-supported clades (Clades 1 – 5; Figure 1; Table 4): Clade 1 is entirely New World in distribution with the exception of a few tramp / invasive species; Clade 2 has an Malagasy / Indo-Pacific distribution; Clade 3 is mostly Old World in distribution; Clade 4 is an entirely Australasian clade; Clade 5 is sister to all remaining *Solenopsis* samples and it contains samples from Australasia and the Neotropics.

**Table 4:** Node support for *Solenopsis* phylogenetic analyses. Note that the BI relaxed clock analysis enforced ingroup monophyly, which is why the root support is not given.

| **Node** | **aLRT** | **aBayes** | **bootstraps** | **PP**<br>**[time-free]** | **PP**<br>**[relaxed**<br>**clock]** |
| --- | --- | --- | --- | --- | --- |
| Clade 1 | 49.1 | 0.940 | 95 | 97.5 | 100 |
| Clade 2 | 100 | 1 | 100 | 100 | 96 |
| Clade 3 | 99.7 | 1 | 100 | 100 | 100 |
| Clade 4 | 100 | 1 | 100 | 100 | 100 |
| Clade 5 | 98.9 | 1 | 100 | 100 | 100 |
| (1,2) | 99.1 | 1 | 100 | 100 | 100 |
| (1,2,3) | 98.5 | 1 | 100 | 100 | 100 |
| (1,2,3,4) | 73 | 0.954 | 78 | 92.8 | 87.7 |
| Root<br>(12345) | 97.8 | 1 | 79 | 79.6 | NA |

**Figure 1.**
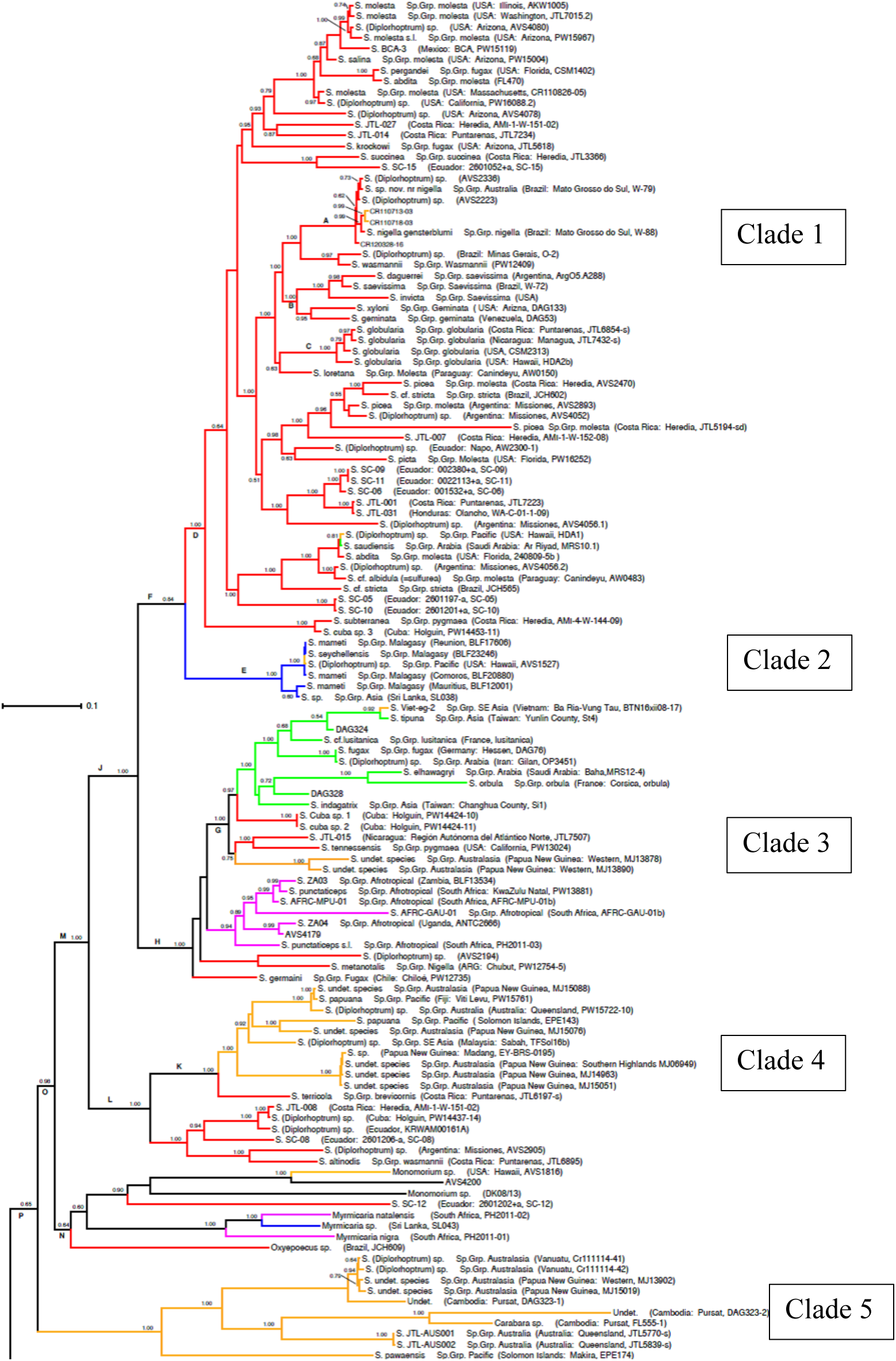
Maximum likelihood phylogeny of *Solenopsis* with branch support values shown (above branch: posterior probability / ultra-fast bootstrap; below branch: SH-aLRT / aBayes). The Bayesian phylogeny, although less resoved in poorly supported areas of the ML tree, is fully congruent with the ML tree. The five numbered major clades correspond to Table 4.

#### Convergence of Bayesian analyses

While the standard diagnostic criteria offered by *MrBayes* and *Tracer* (ESS, ASDSF, PSRF, lnL plots) suggested convergence of 5,000 post-burnin samples from the MCMC (*MrBayes*: ESS[all parameters] ≥ 2,100, ASDSF = 0.02, MSDSF = 0.26, PSRF = [1.000-1.002]; *Tracer*: plateauing of LnL [ESS ≥ 3,700] and peaked marginal posterior probability distributions), more nuanced analyses of convergence implemented in *RWTY* (jumpy Topological Autocorrelation Plots and TreeSpace Plots) suggest that runs where not sampling the same regions of tree space equally. Some metrics calculated by *CODA* confirmed several issues of convergence within and between runs. While the Geweke (1992) Z-score was high (Z > |2.0|) for several parameters, PSRF and multivariate PSRF approached 1.0, Raftery & Lewis’ metric (1992) suggest that for all but two variables sufficient number of iterations were retained and dependence factors (I; the extent to which autocorrelation inflates the required sample size) were close to 1.0 (suggesting there was no undue autocorrelation), Heidelberger & Welch’s Stationarity and Interval Halfwidth Tests were generally passed.

Visualized of tree space using MDS confirmed that the third run, which also had the lowest LnL, sampled from an additional tree island with marginally lower LnL (Suppl. Material Figure 1). Exclusion of this run from did not alter the topology of the ingroup. It was also apparent that the runs sampled the tree islands with differing intensities.

#### Species complexes

With some notable exceptions, most species complexes were not recovered as monophyletic. The fire ants (*S. geminata* complex sensu Trager 1991) were consistently recovered as monophyletic, albeit with low support (Figure 1). In addition, most of the specimens in the *S. nigella* complex formed a monophyletic clade (Figure 1), but one *nigella* representative, *S. metanotalis*, was placed in clade 3. There was very low sequence diversity among the two sampled species of the monophyletic *S. nigella* complex and it is possible that they are representatives of a single species. The *S. wasmanni* complex was not recovered as monophyletic, but three unrelated and well-supported clades emerged: the large-bodied nominal *S. wasmanni*, a Carebarella/*S. altinodis*/*S. iheringi* clade (in clade 1), and a *S. succinea* clade. All other species complexes were never monophyletic clades and no substructure was discernable.

#### Biogeographic Patterns

The phylogeny of *Solenopsis* shows distinct biographic patterns. The large clade 1 is almost entirely New World and represents approximately half of the sampled specimens. Further Neotropical specimens are scattered throughout most other clades. Clade 2 mostly contains species from the Madagascan realm (with singletons from the Oriental [Seychelles Islands] and Nearctic [Hawaii] realms). Clade 3 has a global distribution and contains an Afrotropical clade, a small Oceanian clade, and a Eurasian clade (Palearctic, Oriental, and Saharo-Arabian realms), and a few Neotropical species. There are two Australasian clades (Australian, Oriental, and Oceanian realms; clade 4 and part of clade 5), which are both located at the base of the tree.

The results of S-DIVA and BBM analyses were for very similar to each other (Fig 3; Table 5). *Solenopsis* most likely originated and diversified in the Neotropics. Most dispersal events involve movement from the Neotropics to the Panamanian and Nearctic realms. From the Neotropics, *Solenopsis* also dispersed to the Old World on several occasions. Notably, dispersal to Australasia has occurred twice very early in the evolution of the genus. Since the two basal nodes are Oceanian, this realm always has a non-trivial probability of being ancestral. However, the Neotropical realms (Neoptropics [E] and Panamanian [I]) are always part of the highest probability ancestral areas, thus supporting a neotropical origin for *Solenopsis* (BMM: E (Neotropics) = 0.559, equivocal = 0.18, EI (Neotropics, Panamanian) = 0.084, EF (Neoptropics, Oceanian) = 0.079, and hidden ranges = 0.096) (S-DIVA: E = 0.334, FGI = 0.229, EFGI = 0.224, hidden ranges = 0.212). Without outgroups the inference becomes more vague and always encompasses four realms, but always including the Neotropics (S-DIVA: EFGI (Neotropics, Oceania, Oriental, Panamanian) = 59.63 BEFG (Australian, Neotropics, Oceania, Oriental) = 12.13).

**Table 5.** Historical range reconstructions by statistical divergence with vicariance analysis (S-DIVA) and binary Bayesian method (BMM). Clade letters correspond to labels in Figure 1. A = Australasia, Pacific, & SE Asia, B = North and South America, C = Eurasia, D = Africa, and E = Indian Ocean islands and S Asia

| Clade | S-DIVA | BMM | Description |
| --- | --- | --- | --- |
| A | P(B)=1.00 | P(B)=1.00 | Species Group *nigella* |
| B | P(B)=1.00 | P(B)=1.00 | Fire Ants |
| C | P(B)=1.00 | P(B)=1.00 | Species Group *globularia* |
| D | P(B)=1.00 | P(B)=1.00 | New World Expansion |
| E | P(E)=0.94, P(BE)=0.057 | P(E)=0.82, P(BE)=0.13, P(B)=0.05 | Indian Ocean Clade |
| F | P(BE)=1.00 | P(B)=0.97 |  |
| G | P(B)=0.89, P(AB)=0.10 | P(B)=0.97 | Eurasian Clade + basal lineages |
| H | P(B)=0.94, P(BC)=0.061 | P(B)=1.00 | Clade including Eurasian and African expansions + basal lineages |
| J | P(B)=1.00 | P(B)=0.99 |  |
| K | P(AB)=1.00 | P(B)=0.95 | Australasian clade + *S. terricola* |
| L | P(B)=1.00 | P(B)=0.99 |  |
| M | P(B)=1.00 | P(B)=0.99 | Core *Solenopsis* in both ML and Bayesian |
| N | P(B)=0.66, P(BC)=0.34 | P(B)=0.97 | Putative outgroup clade |
| O | P(B)=0.72, P(BC)=0.28 | P(B)=0.96 | Core *Solenopsis* + putative outgroups (not present in ML) |
| P | P(AB)=0.56, P(ABC)=0.25, P(AC)=0.18 | P(A)=0.47, P(B)=0.46 | All *Solenopsis* with Australasian clade basal (not present in ML) |

**Figure 3.**
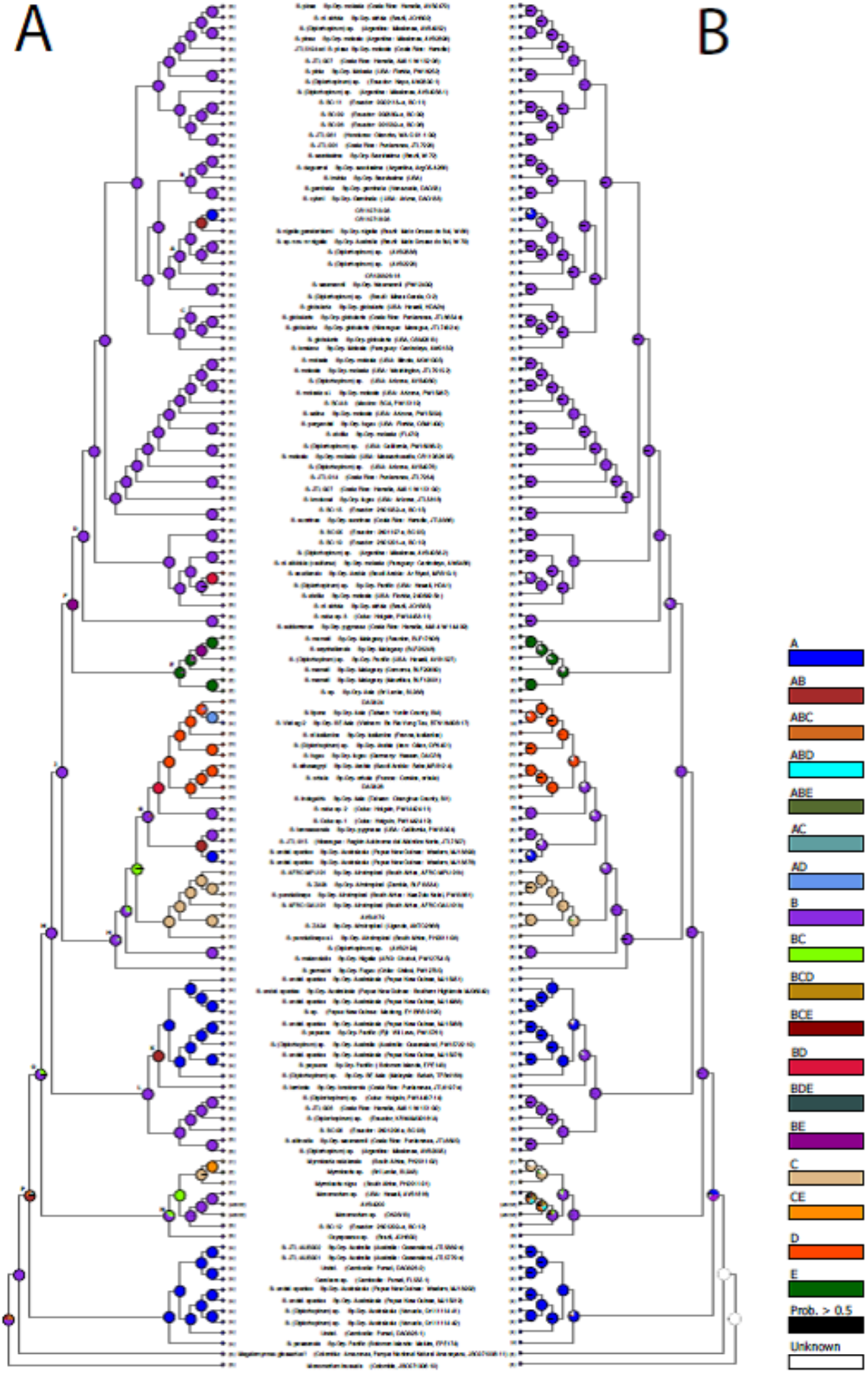
Cladogram showing ancestral range reconstruction of *Solenopsis* based on (A) S-DIVA and (B) Bayesian binary MCMC. Biogeographic ranges are: A = Australasia, Pacific, & SE Asia, B = North and South America, C = Eurasia, D = Africa, and E = Indian Ocean islands and S Asia. Clade labels correspond to Figure 1 and Table 4.

## Discussion

Our analyses consistently converged on the same general topology, although overall branch support was greatly dependent on the method used and some areas where never recovered with high support. In particular, support for the outgroups and the deepest split in *Solenopsis* was always low, regardless of branch support measure. However, the branching order within Solenopsis is identical to that found by Ward et al. (2015), who used many more loci but sampled many fewer species, and highly consistent regardless of tree estimation method.

There was a general trend of high posterior probability and low bootstrap support across many branches on the *Solenopsis* phylogeny. This trend was present at both deep, internal branches and at terminal branches. Disparity between bootstrap values and posterior probabilities is a well-recognized phenomenon (Erixon et al. 2003), despite the fact that they are both interpreted as measures of support for clades in a phylogenetic tree. While posterior probabilities and bootstrap values often do not correspond to each other, there is a positive correlation between them. The strength of the correlation, however, varies from study to study, but bootstrap values are generally lower (Douady et al. 2003). The definition of posterior probability support values as the probability of the tree or a clade given the data assumes the correct model is used, and so the accuracy of Bayesian posterior probabilities more sensitive to model misspecification relative to bootstraps (Huelsenbeck & Rannala 2004). Elevated posterior probabilities are particularly likely if the substitution model is under parameterized, and can lead to strong support for incorrect topologies (Erixon et al. 2003; Huelsenbeck & Rannala 2004). Our MrBayes analysis modeled four of the five genes with GTR+I+Γ and fifth with GTR+Γ, so it seems unlikely that under parameterized models are an issue. However, a site-specific model such as CAT (Lartillot & Philippe 2004; Le et al. 2008) may reveal that long-branch attraction is pulling the putative outgroup clade into *Solenopsis* (Lartillot et al. 2006).

### Taxonomic considerations

It is clear that the genus *Solenopsis* is in need of a comprehensive global revision. The systematic classification of Pacheco & Mackay (2013), which was erected to ease taxonomic identification and not to reflect evolutionary patterns, unsurprisingly bears little resemblance to the molecular phylogenetic relationships recovered in our analysis. However, there are notable some patterns although we emphasize that without complete taxon sampling, caution is due. First, few species complexes are recovered as monophyletic, but there are exceptions. The fire ants (*S. geminata* species complex) are monophyletic. This is not surprising, considering how much attention they have received, relative to their neglected congeners. In addition to the fire ants, only the *S. nigella* and *S. globularia* species complexes were (largely) monophyletic, although this could be an artifact of limited taxon sampling and expanded sampling could erase this pattern. Only the *S. metanotalis* subcomplex of the *S. nigella* complex is an exception and appears quite removed from the *S. nigella* subcomplex. The *S. wasmanni* complex is paraphyletic, but three subcomplexes appear monophyletic: the large and polymorphic *S. wasmanni, S. succinea* (former *Diagyne*), and the remaining species of the complex (the former *Carebarella, S. altinodis*, and *S. iheringi*). Second, several species complexes are restricted to the New World Expansion (clade E), despite not necessarily being monophyletic: *S. geminata, S. globularia, S. molesta*, and *S. stricta*.

In addition, it seems likely that several species may need to be synonymized following careful taxonomic analysis. For example, *S. nigella* and *S. gensterblumi* may be conspecific given the low sequence divergence between the two species. Another species in need of closer taxonomic scrutiny is *S. mameti* and other *Solenopsis* from islands in the Indian Ocean (clade D; Fig 1 & 2). The sample of *S. mameti* from Reunion showed almost no divergence from the conspecific sample from Comoros, on the other side of Madagascar, but moderate divergence from the sample collected on nearby Mauritius (1.5% sequence divergence at shared genes in the dataset). In addition, *S. seychellensis*, from the Seychelles, and an unidentified species from Hawaii also have nearly 100% sequence identity with the Comoros and Reunion *S. mameti*. Population genetic approaches to assess gene flow and genetic differentiation among species in this low divergence clades may be necessary to distinguish widely distributed invasive species from cases of incipient speciation of isolated populations (Smith and Fisher 2009). Yet another noteworthy case is the recently described *S. saudiensis* (Sharaf & Aldawood 2011) from the Arabian Peninsula. It may be synonymous with *S. abdita*, since it is embedded within the New World expansion with strong support and very low sequence divergence among *S. saudiensis*, an unidentified *Solenopsis* collected from Hawaii, and *S. abdita* (Fig 1-2). These samples may well represent a previously unrecognized widespread tramp species since both the authors only considered the regional faunas in their descriptions (Florida for *S. abdita* [Thomas CITE], Saudi Arabia for *S. saudiensis* [Sharaf & Aldawood 2011]). Given the propensity for *Solenopsis* species to be transported by humans, these results imply that new species descriptions should ideally be conducted in the context of global revisions – a truly horrifying notion.

### Biogeography

There is a clear signal of biogeographic structure within *Solenopsis*. Most large clades are associated with particular biogeographic realms, and there are relatively few examples of terminal sister taxa from different biogeographic regions. However, our biogeographic realms are broadly defined, and there is little evidence of finer-scale biogeographic structure within clades. The Indian Ocean, Afrotropics, and Eurasia are all represented by a single clade, suggesting single colonization events with subsequent dispersal and diversification in each region. Dispersal events in the Old World always involve movement to neighboring realms. There is evidence that most, if not all, of these colonization events originated in North or South America, as New World lineages are often found at the base of the expansions into other biogeographic regions. Given the scale of the biogeographic regions used in this study, the associated vicariant events would also have to be on a continental scale. There are a limited number of geological events that could give rise to vicariance on this scale, and vicariance in multiple lineages resulting from a single event would require some degree of genetic or geographic structure already present within the range of the most recent common ancestor of *Solenopsis* ants. Alternatively, multiple dispersal events out of North and South America could explain the pattern, and is supported by the BBM range reconstructions.

The dispersal ability of *Solenopsis* ants has only been studied in a small number of species, mostly in the red imported fire ant *S. invicta*. In the absence of wind, energetic studies of fire ants suggest that dispersal flights after mating are limited to approximately 5 km (Vogt et al. 2000). However, reproductive *S. invicta* have been collected at altitudes as high as 136 m during nuptial flights (Fritz et al. 2011), where winds may aid in long distance dispersal of newly-mated queens. The ability of *S. invicta* and other invasive ants to colonize new habitats is hence not limited to their innate flight capacity.

The ancestral range of the *Solenopsis* genus remains unclear, in part due the uncertainty of the relationships of the basal clades in the *Solenopsis* phylogeny. Taken collectively, the S-DIVA and BBM reconstructions indicate an inability to distinguish Australasia or the New World, or both, as the range of the *Solenopsis* common ancestor. Given the geological association of Australia and South America and the high diversity of *Solenopsis* in South America, it is probable that the New World biogeographic region defined in this study corresponds to South America in the early evolutionary history of the genus. However, the intermingling of North and South American samples on the phylogenetic tree, especially Central and South American samples, makes it nearly impossible to disentangle them analytically. This also suggests a high rate of exchange between to two continents, either after the closure of the Isthmus of Panama and the Central American Seaway approximately three million years ago (Cody et al. 2010), or due to human-mediated movement of species.

## Notes

### Competing Interest Statement

The authors have declared no competing interest.

## References

Ali SS, Yu Y, Pfosser M, Wetschnig W (2012) Inferences of biogeographical histories within subfamily Hyacinthoideae using S-DIVA and Bayesian binary MCMC analysis implemented in RASP (Reconstruct Ancestral State in Phylogenies). Annals of Botany, 109, 95–107.

Anisimova M, Gil M, Dufayard JF, Dessimoz C, Gascuel O (2011) Survey of branch support methods demonstrates accuracy, power, and robustness of fast likelihood-based approximation schemes. Systematic Biology, 60, 685–699.

Brown JM (2014a) Predictive approaches to assessing the fit of evolutionary models. Systematic Biology, 63, 289–292.

Brown JM (2014b) Detection of implausible phylogenetic inferences using posterior predictive assessment of model fit. Systematic Biology, 63, 334–348.

Cody S, Richardson JE, Rull V, Ellis C, Pennington RT (2010) The great American biotic interchange revisited. Ecography, 33, 326–332.

DeHeer CJ, Goodisman MAD, Ross KG (1999) Queen dispersal strategies in the multiple queen form of the fire ant *Solenopsis invicta*. American Naturalist, 153, 660–675.

Douady CJ, Delsuc F, Boucher Y, Doolittle WF, Douzery EJP (2003) Comparison of Bayesian and maximum likelihood bootstrap measures of phylogenetic reliability. Molecular and Biological Evolution, 20, 248–254.

Erixon P, Svennblad B, Britton T, Oxelman B (2003) Reliability of Bayesian posterior probabilities and bootstrap frequencies in phylogenetics. Systematic Biology, 52, 665–673.

Folgarait PJ (1998) Ant biodiversity and its relationship to ecosystem functioning: A review. Biodiversity and Conservation, 7, 1221–1244.

Fritz GN, Fritz AH, Vander Meer RK (2011) Sampling high-altitude and stratified mating flights of red imported fire ant. Journal of Medical Entomology, 47, 508–512.

Guindon S, Dufayard JF, Lefort V, Anisimova M, Hordijk W, Gascuel O (2010) New Algorithms and Methods to Estimate Maximum-Likelihood Phylogenies: Assessing the Performance of PhyML 3.0. Systematic Biology, 59, 307–321.

Holt BG, Lessard J-P, Borregaard MK, Fritz SA, Araújo MB, Dimitrov D, Fabre P-H, Graham CH, Graves GR, Jønsson KA, Nogués-Bravo D, Wang Z, Whittaker RJ, Fjeldså J, Rahbek C (2013) An Update of Wallace’s Zoogeographic Regions of the World. Science, 339, 74–78.

Huelsenbeck JP, Rannala B (2004) Frequentist properties of Bayesian posterior probabilities of phylogenetic trees under simple and complex substitution models. Systematic Biology, 53, 904–913.

Huelsenbeck JP, Ronquist F (2001) MRBAYES: Bayesian inference of phylogeny. Bioinformatics, 17, 754–755.

Jombart T, Kendall M, Almagro-Garcia J, Colijn C (2017) treespace: Statistical exploration of landscapes of phylogenetic trees. Molecular Ecology Resources, DOI: 10.1111/1755-0998.12676.

King JR, Tschinkel WR, Ross KG (2009) A case study of human exacerbation of the invasive species problem: transport and establishment of polygyne fire ants in Tallahassee, Florida, USA. Biological Invasions, 11, 373–377.

Lartillot N, Brinkmann H, Philippe H (2007) Suppression of long-branch attraction artefacts in the animal phylogeny using a site-heterogeneous model. BMC Evolutionary Biology, 7, article S4.

Lartillot N, Philippe H (2004) A Bayesian mixture model for across-site heterogeneities in the amino-acid replacement process. Molecular Biology and Evolution, 21, 1095–1109.

Le SQ, Gascuel O, Lartillot N (2008) Empirical profile mixture models for phylogenetic reconstruction. Bioinformatics, 24, 2317–2323.

Liu L (2008) BEST: Bayesian estimation of species trees under the coalescent model. Bioinformatics, 24, 2542–2543.

Liu L, Pearl DK (2007) Species trees from gene trees: reconstructing Bayesian posterior distributions of a species phylogeny using estimated gene tree distributions. Systematic Biology, 56, 504–514.

McGlynn TP (1999) The worldwide transfer of ants: Geographical distribution and ecological invasions. Journal of Biogeography, 26, 535–548.

Moser JC, Blomquist SR (2011) Phoretic arthropods of the red imported fire ant in central Louisiana. Annals of the Entomological Society of America, 104, 886–894.

Nylander JAA, Olsson U, Alström P, Sanmartín I (2008b) Accounting for phylogenetic uncertainty in biogeography: a Bayesian approach to Dispersalvicariance Analysis of the thrushes (Aves: Turdus). Systematic Biology, 57, 257–268.

O’Meara, BC (2010) New Heuristic Methods for Joint Species Delimitation and Species Tree Inference. Systematic Biology, 59, 59–73.

Pacheco JA (2007) The New World thief ants of the genus *Solenopsis* (Hymenoptera: Formicidae). Doctoral Dissertation. University of Texas, El Paso, Texas.

Passera L (1994) Characteristics of tramp species. In Williams DF Exotic Ants: Biology, Impact, and Control of Introduced Species, pp. 23–43. Boulder, CO: Westview.

Pitts JT, McHugh JV, Ross KG (2005) Cladistic analysis of the fire ants of the *Solenopsis saevissima* species-group (Hymenoptera: Formacidae). Zoologica Scripta, 34, 493–505.

Plummer M, Best N, Cowles K, Vines K (2006). CODA: Convergence Diagnosis and Output Analysis for MCMC, R News, vol 6, 7–11.

Poldi B (1992) A poorly known taxon: *Diplorhoptrum orbulum* (Emery 1875). Ethology, Ecology, and Evolution, 2, 91–94.

Powell et al. (2020) Frontiers in Genetics

R Core Team (2017) R: A language and environment for statistical computing. R Foundation for Statistical Computing, Vienna, Austria. URL https://www.R-project.org/.

Ronquist F (1997) Dispersal-vicariance analysis: A new approach to the quantification of historical biogeography. Systematic Biology, 46, 195–203

Ronquist F, Huelsenbeck JP (2003) MRBAYES 3: Bayesian phylogenetic inference under mixed models. Bioinformatics, 19, 1572–1574.

Ross KG, Gotzek D, Ascunce MS, Shoemaker DD (2010) Species delimitation: A case study in a problematic ant taxon. Systematic Biology, 59, 162–184.

Ross KG, Krieger MJB, Keller L, Shoemaker DD (2007) Genetic variation an dstructure in native populations of the fire ant *Solenopsis invicta*: evolutionary and demographic implications. Biological Journal of the Linnean Society, 92, 541–560.

Ross KG, Shoemaker DD (2005) Species delimitation in native South American fire ants. Molecular Ecology, 14, 3419–3438.

Sharaf MR, Aldawood AS (2011) First occurrence of *Solenopsis* Westwood 1840 (Hymenoptera: Formicidae), in the kingdom of Saudi Arabia, with description of a new species *S. saudiensis* n. sp. Annales de la Société Entomologique de France, 47, 474–479.

Schenk et al. 2016 PLoS One

Smith MA, Fisher BL (2009) Invasions, DNA barcodes and rapid biodiversity assessment using ants of Mauritius. Frontiers in Zoology, 6, 31.

Vogt, JT, Appel AG, West MS (2000) Flight energetics and dispersal capability of the fire ant, *Solenopsis invicta* Buren. Journal of Insect Physiology, 46, 697–707.

Yang CC, Ascunce MS, Luo LZ, Shao JG, Shih CJ, Shoemaker D (2012) Propagule pressure and colony social organization are associated with the successful invasion and rapid range expansion of fire ants in China. Molecular Ecology, 21, 817–833.

Yu Y, Harris AJ, Blair C, He XJ (2015) RASP (Reconstruct Ancestral State in Phylogenies): a tool for historical biogeography. Molecular Phylogenetics and Evolution, 87, 46–49.

Warren D, Geneva A, Swofford D, Lanfear R (2017) RWTY (R We There Yet): An R Package for Examining Convergence of Bayesian Phylogenetic Analyses. Molecular Biology and Evolution, 34, 1016–1020.

